# A simple canonical circuitry for non-stationary normalization by inhibition to explain and predict change detection in monkey area MT

**DOI:** 10.1101/2020.06.19.161414

**Authors:** Udo Ernst, Xiao Chen, Lisa Bohnenkamp, Fingal Orlando Galashan, Detlef Wegener

## Abstract

Sudden changes in visual scenes often indicate important events for behavior. For their quick and reliable detection, the brain must be capable to process these changes as independent as possible from its current activation state. In motion-selective area MT, neurons respond to instantaneous speed changes with pronounced transients, often far exceeding the expected response as derived from their speed tuning profile. We here show that this complex, non-linear behavior emerges from the combined temporal dynamics of excitation and divisive inhibition, and provide a comprehensive formal analysis. A central prediction derived from this investigation is that attention increases the steepness of the transient response irrespective of the activation state prior to a stimulus change, and irrespective of the sign of the change. Extracellular recordings of attention-dependent representation of both speed increments and decrements confirmed this prediction and suggest that improved change detection derives from basic computations in a canonical cortical circuitry.

## Introduction

For change detection in behaviorally relevant situations, stimulus-induced, strong transient firing rate modulations provide a powerful signal to downstream visuomotor areas on short timescales^1,2^. Such transients are particularly information-rich^3,4^, and the temporal sensitivity of neurons to stimulus changes is suggested to constitute a major function of neuronal tuning properties^5^. Human behavioral performance to detect speed changes^6^ correlates with the size of area MT change transients^7^ and strongly improves with both spatial and feature-directed attention^8^. Accordingly, both forms of attention were found to increase sustained and transient MT firing rates before and after a stimulus change^9,10^, and the transient response after the change also correlates with reaction times^9^.

This close relation between neuronal transients and behavioral change detection performance on the one hand, and the general effect of attention to increase neuronal response rates^11,12^ on the other raises the question how attention exerts its beneficial effects under conditions where firing rate increases seem to impede the formation of pronounced transients and counteract behavioral performance. For example, because attention-dependent enhancement of firing rates brings a neuron closer to its maximum activity, a stimulus-induced transient firing rate increase under attention might be smaller than without attention. Furthermore, a negative transient would start from a larger pre-change activation level and is presumably having a smaller absolute negative peak than under conditions of no or remote attention. Absolute firing rates of the transient, therefore, may only poorly allow to predict attention-dependent improvements of behavioral change detection. As a consequence, the neuronal circuit processing the change is supposed to rely on some form of normalization to compensate for differences in absolute activity, and, to facilitate change detection, attentional modulation of the circuit should induce a consistent effect on neuronal transients that is independent from the stimulus-induced activation level before the change. It is unclear presently, which feature of a change-transient can be most consistently modulated by attention independent of the specific stimulus condition, and which neuronal mechanism might underlie its corresponding dynamics.

To investigate this issue, we set up and formally analyze a biologically plausible, canonical circuit providing divisive inhibition to an excitatory unit, and apply the model to a wide range of different stimulus conditions under passive viewing conditions^7^. By introducing an input response gain to simulate top-down modulation of change detection by visual attention, the model circuit predicts a main effect on response rise times not only for positive transients but also for the case of negative transients, i.e. rapid rate decreases in a population of neurons. To test this prediction experimentally, we recorded single neurons from area MT while monkeys were engaged in a change detection task for both speed increments and decrements, eliciting large positive and negative population transients, respectively. We show that MT activity is exactly following the prediction of the model, having steeper response slopes irrespective of the sign of the transient. Thus, the model circuit provides a consistent neuronal mechanism to explain change detection under passive and attended conditions by a rather simple computational unit realized in the canonical circuitry of the cortex.

## Results

### Transient neural responses in area MT

Neurons in area MT are well-activated by moving stimuli such as localized drifting Gabor patches presented inside their receptive fields (RFs). Neural responses are tuned to the direction of movement, exhibiting maximal activity when the stimulus is moving in the cell’s preferred direction^13^. The additional preference of MT neurons for a particular stimulus speed can be approached by a Gaussian function in log-velocity space^14,15^. Rapid changes in the input to these neurons result in pronounced, transient changes of their activation. An example from previous work^7^ is showing the population response of an ensemble of MT neurons to a range of sudden stimulus accelerations and decelerations (Fig. 1a). Transients show up as fast increases (or decreases) in firing rate, followed by a slower decrease (or increase) in firing rate back to a sustained level of activation. Such transients are visible in response to any instantaneous change in stimulation (black arrows in Fig. 1a): stimulus onset, motion onset, speed change, motion offset, and stimulus offset. Due to their causal relation to visual perception and change detection^16^, we aimed to study their properties and non-linear dynamics in a theoretical framework providing access to a comprehensive formal analysis, and to experimentally test predictions derived from it.

**Figure 1.**
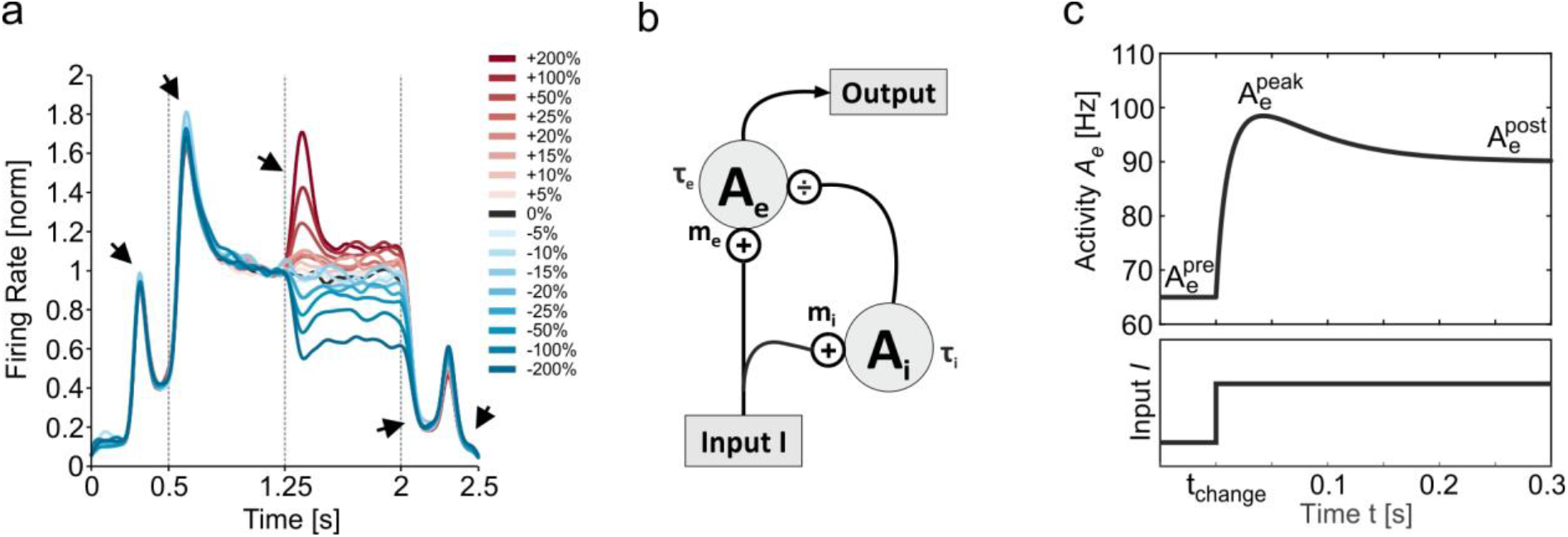
Transient responses in area MT and model circuit. **(a)** Population response of MT neurons exhibiting large transients (black arrows) in response to stimulus onset (at t = 0.25 s), movement onset (at t = 0.5 s), speed change (at t = 1.25 s), movement offset (at t = 2.0 s) and stimulus offset (at t = 2.25 s). Color scale indicates sign and magnitude of the speed change (blue for decelerations, red for accelerations). Modified from Ref. [7]. **(b)** Model circuit providing divisive inhibition Ą to an MT neuron with activation *A_e_. τ_e_* and *τ_i_* denote time constants and *m_e_* and *m_i_* denote gains of the corresponding units. **(c)** Example response of the model circuit (upper graph) to a positive input change at time *t_change_* (lower graph). Starting from activation level 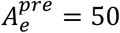 spikes/s, the circuit exhibits a fast transient increase in activation reaching a peak response 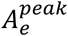 before decreasing more slowly to a post-change, sustained activation level 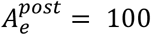 spikes/s (with parameters: *τ_e_* = 10 ms, *τ_i_* = 40 ms, 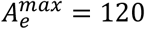 spikes/s).

### Model for transient activation in area MT

For modeling transient responses, we consider a circuit where an input *I* activates an excitatory and an inhibitory unit, with output *A_i_* of the inhibitory unit providing divisive inhibition onto the excitatory unit (Fig. 1b). The dynamics of the circuit is given by the following differential equations:

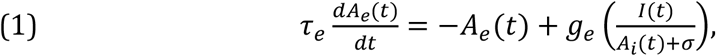

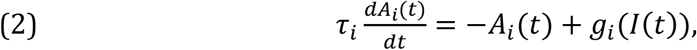

where output *A_e_*(*t*) of the excitatory unit represents the instantaneous firing rate *A_e_* of a neuron in MT at time *t*, with *I* being the external input provided by the visual stimulus. Terms *τ_e_, τ_i_* denote time constants, *σ* is a constant positive offset, and *g_e_, g_i_* represent gain functions realized by threshold-linear rectification, with *m_e_*, *m_i_* indicating gain factors, and *θ_e_, θ_i_* indicating thresholds:

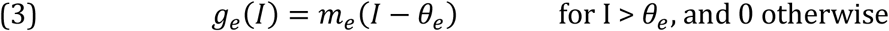

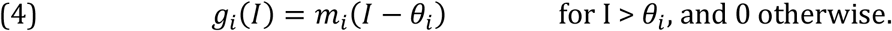

With constant suprathreshold input *I*_0_, the steady-state solution

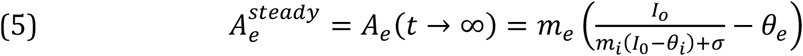

for equation (1) is equivalent to a standard divisive normalization model^17,18^, where *A_i_* represents feedback from co-activated neighboring MT columns. As such, our model can be interpreted as a dynamical reformulation of static divisive normalization. Its advantage is the explicit representation of transients, allowing activation *A_e_* to quickly follow changes in *I* on a fast time scale *τ_e_*, while divisive inhibition is acting on a longer time scale *t_i_*, bringing the output towards a sustained, steady-state level of activation (Fig. 1c).

### Model fit to transients from experimental data

To investigate how well the model explains experimental data, *A_e_*(*t*) was fitted to MT stimulus onset transients. Because MT neurons exhibit spontaneous activity with low firing rates even in absence of a visual stimulus, we assume *θ_e_* = *θ_i_* = 0 and *I*(*t*) > 0, which allows to replace equations (3) and (4) by linear gain functions to simplify the analysis. By modeling stimulus onset as an instantaneous change in external input at *t* = *t_change_* from *I^pre^* to *I^post^*, equation (2) can be explicitly solved, assuming that for *t* < *t_change_* the system is in its steady state for a constant input *I^pre^*. The result can be rewritten in terms of the sustained activation 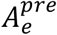 before stimulus onset, and sustained activation 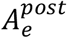 after decay of the transient response, to account for the fact that experimental access is given to the output of the neuron rather than to its input:

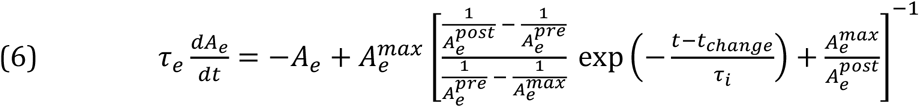

The ratio of the two gain factors 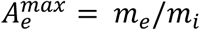 designates the theoretical maximum sustained activation of the circuit. We used a grid search to find the remaining free parameters *τ_e_*, *τ_i_*, and 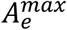 to minimize the average quadratic error between model activation and recorded MT firing rate during the transient response (cf. Methods).

This structurally very simple model allowed for a close approximation of transient and sustained MT responses to motion onsets, despite the significant differences in their shape, caused by, among other things, the different tuning of individual neurons to actual stimulus speeds. Three examples of multi-unit responses and corresponding model fits are given in Fig. 2a-c, and the goodness-of-fit distributions for the total of 88 (monkey 1 (M1)) and 55 (M2) units are shown in Figure 2d. The goodness-of-fit ratio G was close to 1 for most units, indicating that fits were estimating the mean response as good as the experimental data (cf. Methods for more details). Excitatory time scales were much faster than inhibitory ones, as to be expected for the dynamics of transients (Fig. 2e). For M1, mean *τ_e_* was 18 ± 16 ms SD, mean *τ_i_* was 67 ± 38 ms, average ratio *τ_i_/τ_e_* was 6.65 with SD 8.56, and average 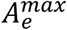 was 70 ± 45 spikes/s. For M2, mean *τ_e_* was 20 ± 10 ms, mean *τ_i_* was 78 ± 34 ms, average ratio *τ_i_/τ_e_* was 5.6 with SD 5.64, and average 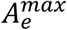 was 128 ± 102 spikes/s.

**Figure 2.**
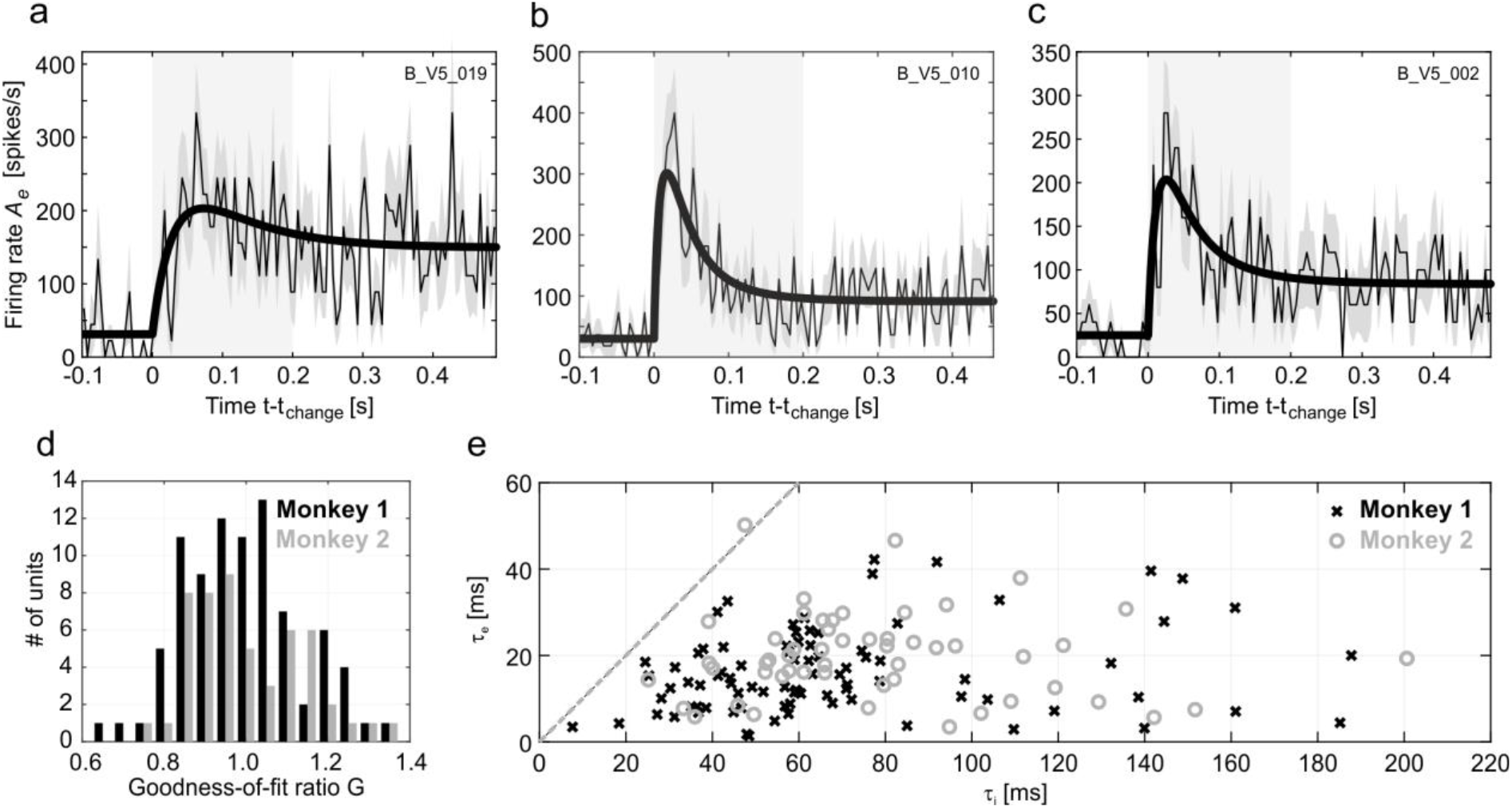
Model fits to stimulus onset transients. **(a – c)** Activity of three example units (thin lines) and corresponding model fit (solid lines). Gray shading indicates standard error over trial repetitions, light gray box indicates fitting interval. Time axis is shown relative to neural response onset. **(d)** Model goodness-of-fit ratio distributions of the recorded units for two monkeys (M1, black; M2, gray). **(e)** Scatter plot for fitted time scales *τ_e_* versus *τ_i_* (M1, crosses; M2, open circles).

### Transient response characteristics to stimulus changes

After the model had been calibrated to motion onset transients, we were next interested to investigate whether it is capable to predict and explain the complex non-linear scaling of transients in response to changes in speed, which depend on the sign and magnitude of the physical speed change and the individual neurons’ tuning characteristics^7^. Because time scales *τ_e_* were shorter than *τ_i_*, one can consider an idealized version of the model by assuming that the excitatory unit reacts infinitely fast to input changes. With this approximation, it is possible to solve equation (6) explicitly for *τ_i_* > 0 and obtain the peak of the transient shortly after *t* = *t_change_* analytically (cf. equations (12) & (13) in Methods).

Peak amplitudes obtained from the model in this manner reproduced an important and so far unexplained characteristic of MT change transients. In MT, peak amplitudes in response to speed changes exceed those expected from the neuron’s speed tuning profile significantly if the speed before the change is away from the neuron’s preferred speed. Neurons that are well tuned to the pre-change speed are very poor change detectors, while neurons for which the pre-change speed is on the flank of their tuning curve have a strong impact on the population response^7^. This tuning-dependent, non-linear relationship between pre-change stimulus speed and individual neuronal tuning profiles is well captured by the model. Simulated speed changes, realized by step functions applied to the circuit’s input at time *t_change_* (Fig. 1c), predict speed change transients very closely matching the experimental data with regard to sign and amplitude of the peak for both changes occurring on the ascending and the descending flank of the tuning curve (Fig. 3). Thus, the model reproduces the full dynamics of physiological change transients, including the over- and undershooting of peak amplitudes.

**Figure 3.**
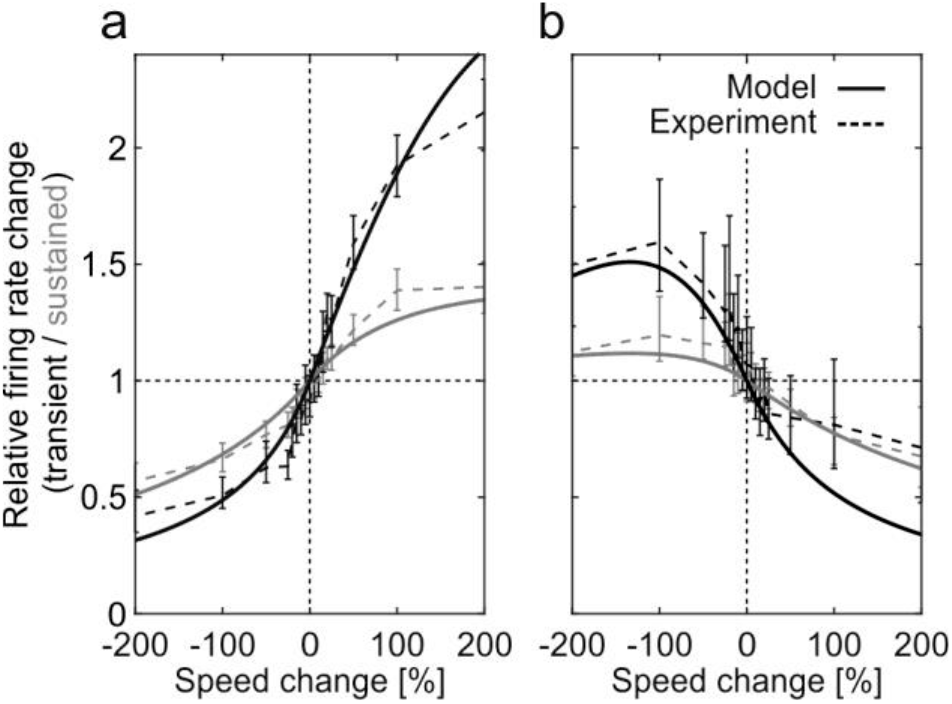
Transient/sustained rate changes following stimulus speed changes. Experimentally observed transient (black) and sustained (gray) rate changes in response to positive and negative speed changes (dashed lines) of different magnitude, and corresponding predictions from model analysis (solid lines). Base speed before the change was located either on **(a)** the ascending flank, or **(b)** the descending flank of the cells’ speed tuning curves.

### Model predictions for attention-dependent modulation of change transients

Because transients signal stimulus changes in a rapid and pronounced manner, they were previously suggested as a possible target for attentional modulation and a neuronal mechanism to speed up reaction times^9,19,20^. In area MT, using speed changes of 100% magnitude, attention was found to modulate both the peak amplitude and latency of a change transient^9,10^. Modulations due to other magnitudes of change, including negative changes, have not been investigated yet. Therefore, in our model, we next studied the dynamical effects of attention on transients for speed changes of arbitrary magnitude. We included attention by simply assuming a multiplicative scaling^11^ of the input *I* by a factor *α* >1, *I* → *αI* (Fig. 4a, top left).

**Figure 4.**
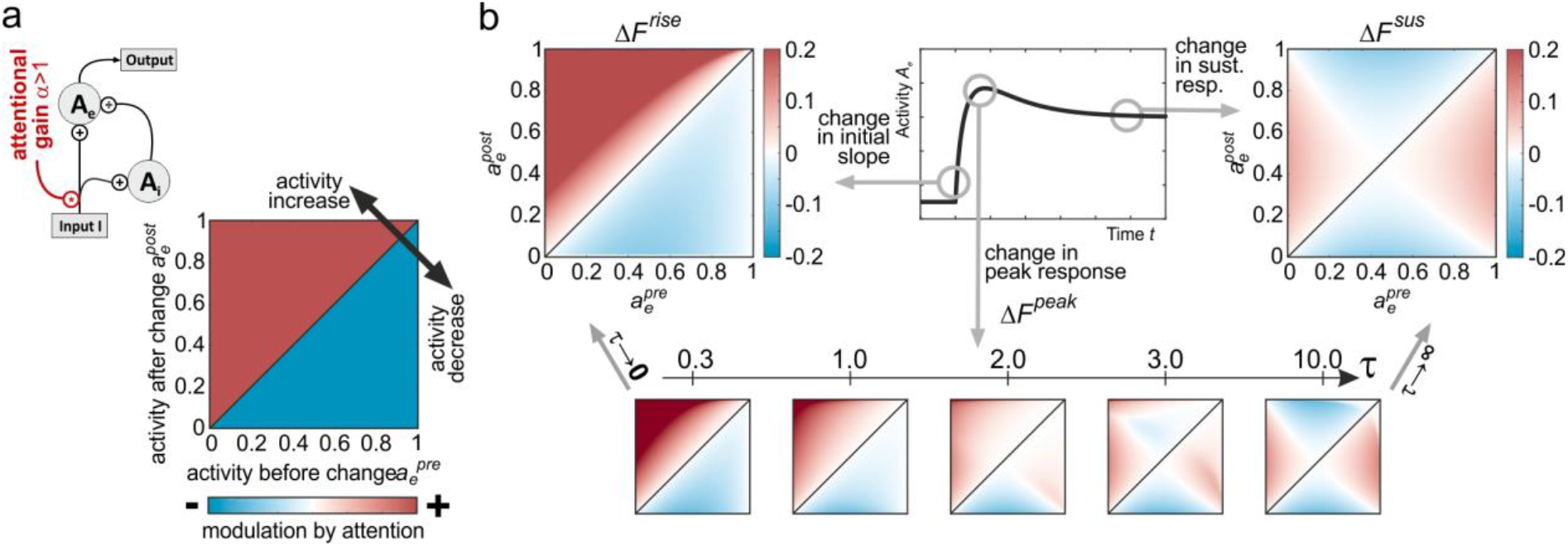
Model predictions for attention-induced changes in slope, peak, and sustained responses. **(a)** Attention is included into the model by a simple multiplicative gain change of the input (top left schematic). Ideally, attention should elicit a consistent modulation of the neuronal change response, as e.g. enhancing a neural response feature for positive input changes (above diagonal, red shading), and suppressing it for negative input changes (below diagonal, blue shading) irrespective of the *particular* pre-change activation level. **(b)** Predicted attention-dependent changes in slope Δ*F^rise^* (top left, indicating consistent modulation by attention), sustained activation *ΔF^sus^* (top right, indicating inconsistent, pre-change activation-dependent modulation by attention), and relative peak height *ΔF^peak^* (bottom plots). Relative peak height critically depends on *τ* and becomes similar to *ΔF^rise^* for *τ* → 0 and similar to *ΔF^sus^* for *τ* → ∞. Color scale for bottom plots is the same as for top plots.

Ideally, to improve computation and, ultimately, perception, the effect of attention should be consistent across the entire dynamical range of the circuit. In particular, for attention to be effective, any change in the external input *I* causing a positive (or negative) transient (as e.g. the traces for the ± 100% changes in Fig. 1a) should be associated with attention-dependent modulations preserving the sign of the transient independent from the neuron’s pre-change activation, as e.g. a consistent increase (or decrease) in the transient’s peak amplitude. If, however, peak amplitude is only increased for low pre-change activation levels, but decreased for high pre-change activation levels for otherwise identical stimulus conditions, the effect of attention would be inconsistent. Figure 4a (bottom right) exemplifies a consistent pattern of attentional modulation for stimulus-change responses associated with positive and negative rate changes, normalized to a dynamic range between 0 (lowest level) and 1 (highest level). The colored surface illustrates a consistent (i.e. pre-change activity independent) positive modulation by attention for stimulus-induced rate increases (above diagonal), and a corresponding negative modulation by attention for stimulus-induced rate decreases (below diagonal).

The model was used to investigate three different change-response features regarding their dependence from attention and pre-change activity: the initial slope *F^rise^* of the transient (i.e. rise/decay time), the maximal amplitude *F^peak^* of the transient, and the sustained activation *F^sus^* following the transient. For analysis, pre- and postchange activities 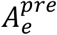 and 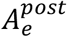 were normalized from absolute values to activations 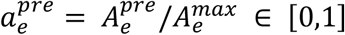 and 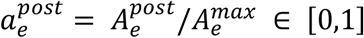 relative to the hypothetical maximum response 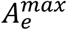. Peak and sustained responses were quantified relative to pre-change activation, assuming that any plausible change detection circuit needs to base its computation on the number of spikes exceeding or falling below this level. The initial slope becomes

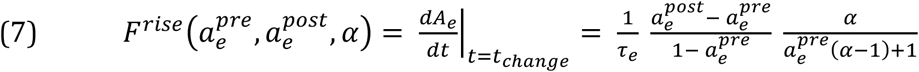

and its attention-induced change Δ*F^rise^* is visualized in Fig. 4b (left). Similarly, the sustained activation level *F^sus^* is given by

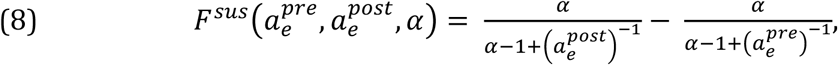

and its attention-induced change Δ*F^sus^* relative to the pre-change activity is plotted in Fig. 4b (right). Finally, the relative change in maximal amplitude Δ*F^peak^* depends on the specific ratio of the excitatory and inhibitory time constants *τ = τ_e_/τ_i_*. Numerical evaluations for different values of *τ* reveal that for *τ* → 0, Δ*F^peak^* approaches Δ*F^rise^*, while for *τ → ∞*, it approaches *ΔF^sus^* (Fig. 4b, bottom panels). These computations reveal two main insights: First, the slope and the sustained response indicate the two extremes of the analysis. The slope is subject to a strong and consistent pattern of attentional modulation, which is independent of both the overall activity and the sign of the rate change, indicating generally faster transients with attention for both speed increments and decrements. In contrast, the sustained response exhibits an inconsistent pattern of attentional modulation, with the sign of the modulation depending on the overall activation of the neuron before the change. Second, as a function of the specific ratio of excitatory and inhibitory time constants, the modulation of the peak amplitude shifts between these extremes. Attentional modulation becomes stronger and more consistent the smaller the ratio *τ = τ_e_/τ_i_*, but is attenuated (and, theoretically, even inconsistent) for larger values of *τ*. Thus, the model makes the explicit prediction that, like positive transients, negative transients (i.e. rapid decreases in firing rate) will have a steeper rise time, i.e. shorter latencies, under the influence of attention as well as higher relative peaks, assuming *τ* being in the previously estimated range (Fig. 2e).

### Experimental investigation of model predictions

Because attention is generally found to *facilitate* the response of visual neurons to the initial stimulus, any *decrease* in the population firing induced by a corresponding change of the behaviorally relevant stimulus would be antagonized by the opposite effect of attention, in terms of absolute firing. For this problem, the model’s prediction of generally faster rise times and potentially more pronounced relative peak firing rate changes as a result of attention offers a particularly attractive solution for effective detection of stimulus changes. Changes in rise time (and, under appropriate conditions, relative peak firing rate changes), provide a mechanism independent of absolute firing to transmit attentionally selected information to downstream areas of the visuomotor pathway.

To test the model predictions, we recorded neuronal responses from motion-sensitive area MT of two macaques (M3: *N* = 45 units, M4: *N* = 25 units). Monkeys performed a speed-change detection task requiring them to either attend towards or away from the recorded unit’s RF and to detect increments or decrements of the target’s speed (Fig. 5a). Peri-stimulus time histograms (PSTHs) aligned to the speed change of the stimulus displayed higher pre-change firing rates when the stimulus inside the RF was attended, and strong, transient increases and decreases of the firing rate in response to increments (accelerations) and decrements (decelerations) of motion speed, respectively (Fig. 5b). Interestingly, attentional modulation before the change did not differ between blocks of speed increments and decrements (two-sided Wilcoxon signed rank test on the attentional modulation index 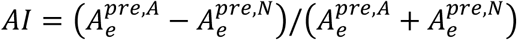, *W = 276, P* = 0.10, *N* = 43 for M3 and *W* = 23, *P* = 0.70, *N* = 21 for M4), indicating that it was independent from the sign of the speed change to be detected. Spike counts for all attentional conditions and all 25 ms time intervals between 400ms before and 200ms after a speed change exhibited a variance close to their mean (Fig. 5b, insets), compatible with the statistical properties of a Poisson process (used below for assessing significance of spike count differences).

**Figure 5.**
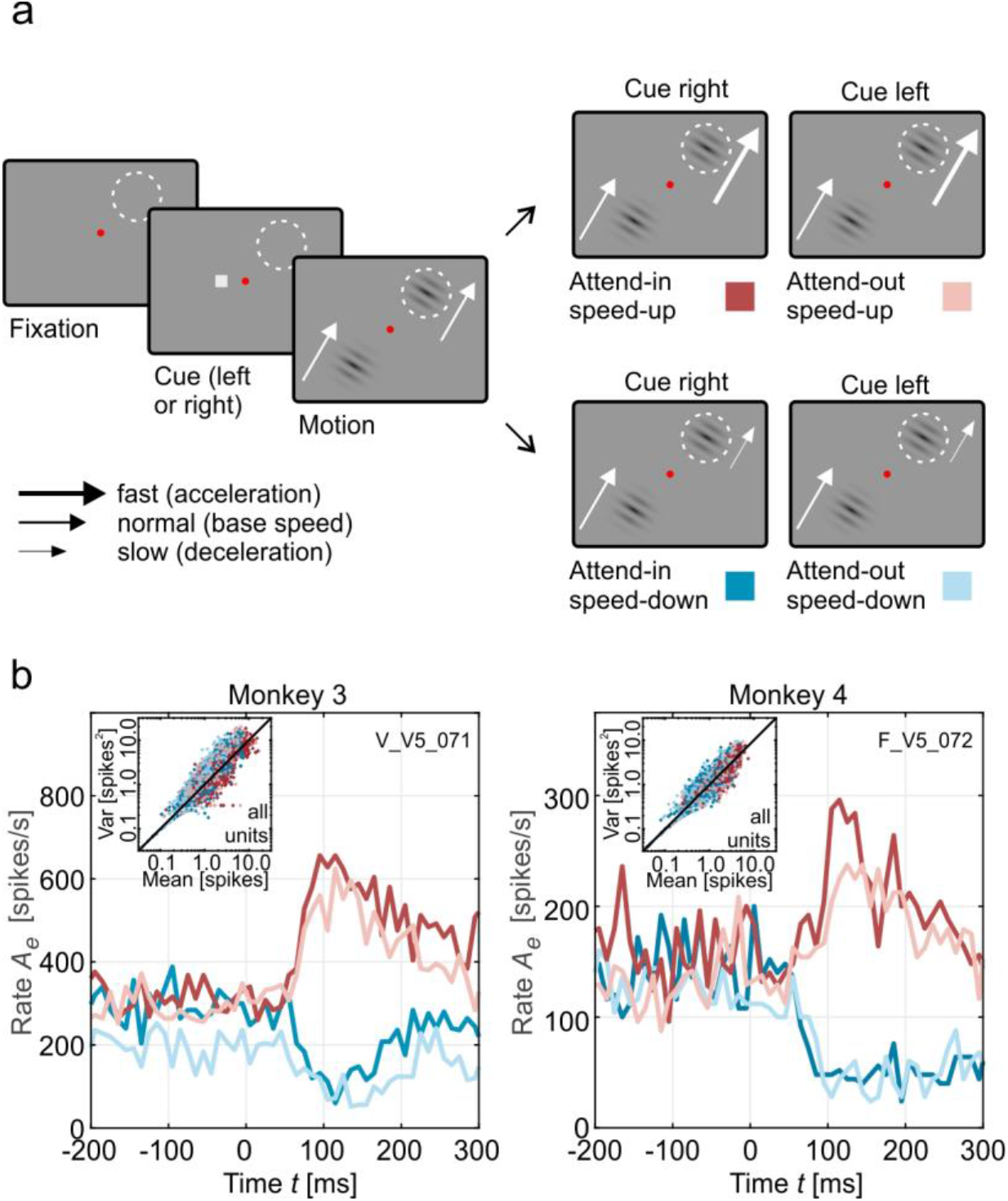
Experimental speed change detection paradigm and example PSTHs. **(a)** Monkeys were detecting positive and negative speed changes (presented block-wise) at pre-cued locations. Each trial started with a cue indicating the hemifield of the rewarded stimulus (small gray box, left or right of the central, red fixation spot). After monkeys properly fixated and pressed a lever, two moving gratings appeared, one of which was placed inside the RF of the recorded neuron (dashed white circle), while the other was placed in the opposite hemifield, mirrored across the fixation spot. Following a pseudo-randomized delay of 0.66 to 5.5 s, the RF-stimulus rapidly increased (top row) or decreased speed (bottom row). Any speed change of the stimulus in the uncued hemifield had to be ignored. Keeping fixation and releasing the lever within 750 ms after the speed change was rewarded with some drops of water of diluted grape juice. Depending on the cued stimulus location, the RF stimulus was either attended or non-attended, giving rise to four experimental conditions (right plots, color-coded). **(b)** PSTHs for two example multi-unit sites (one from each monkey), illustrating rapid firing rate adjustments in response to speed changes. Note that because negative speed changes result in a decrease of the firing rate, the stimulus-induced modulation of the firing rate is opposite to the attention-induced modulation before the change. Insets show spike counts and variances assessed in time intervals of 25 ms from −400ms to +200ms relative to stimulus change for all units. Color coding of the different attentional conditions is indicated in panel (a).

To analyze these transients according to model predictions, we assessed first, response slopes and second, relative firing rate changes during the transient time period of 50 to 200 ms following the speed change. First, for analyzing slopes, we calculated excess cumulative spike counts representing the number of spikes over- or undershooting the mean firing rate before the speed change as a proxy for the initial slope. Because rates increase or decrease almost linearly for the first 40 ms following the population transient onset, a larger (smaller) cumulative count is equivalent to a steeper positive (negative) slope. In contrast to estimating slopes directly from the PSTHs, the cumulative measure has two advantages. First, because accumulation is based on integration, noise does not become amplified but attenuated. Second, accumulation does not need smoothing of data to more reliably calculate specific response parameters, which is problematic if transients are faster than the width of the PSTH filter kernel used. Consistent with the prediction of the model, the excess cumulative spike count was found to rise more steeply for positive transients, and to decay more rapidly for negative transients when attention was directed to the stimulus (Fig 6a, b). Onset of the population transient was around 55 ms after the stimulus change, which is well in the range of typical MT response latencies^21,22^. Attention-dependent differences in excess cumulative spike counts become significantly different from 0 already shortly after response onset (Fig. 6c). We also tested this result at the level of individual units (Figs. 6 d, e). Of all units, 60% displayed significantly different cumulative spike counts during the onset of the transient following speed increments, and of those, 85% confirmed the predictions of the model. Likewise, for speed decrements, 46% of all units were displaying significantly different spike counts, and 76% of those were confirming the model’s prediction of steeper slopes. This result was found for both monkeys, with 84% (M3) and 72% (M4) of all significantly modulated units being in accordance with the model prediction.

**Figure 6.**
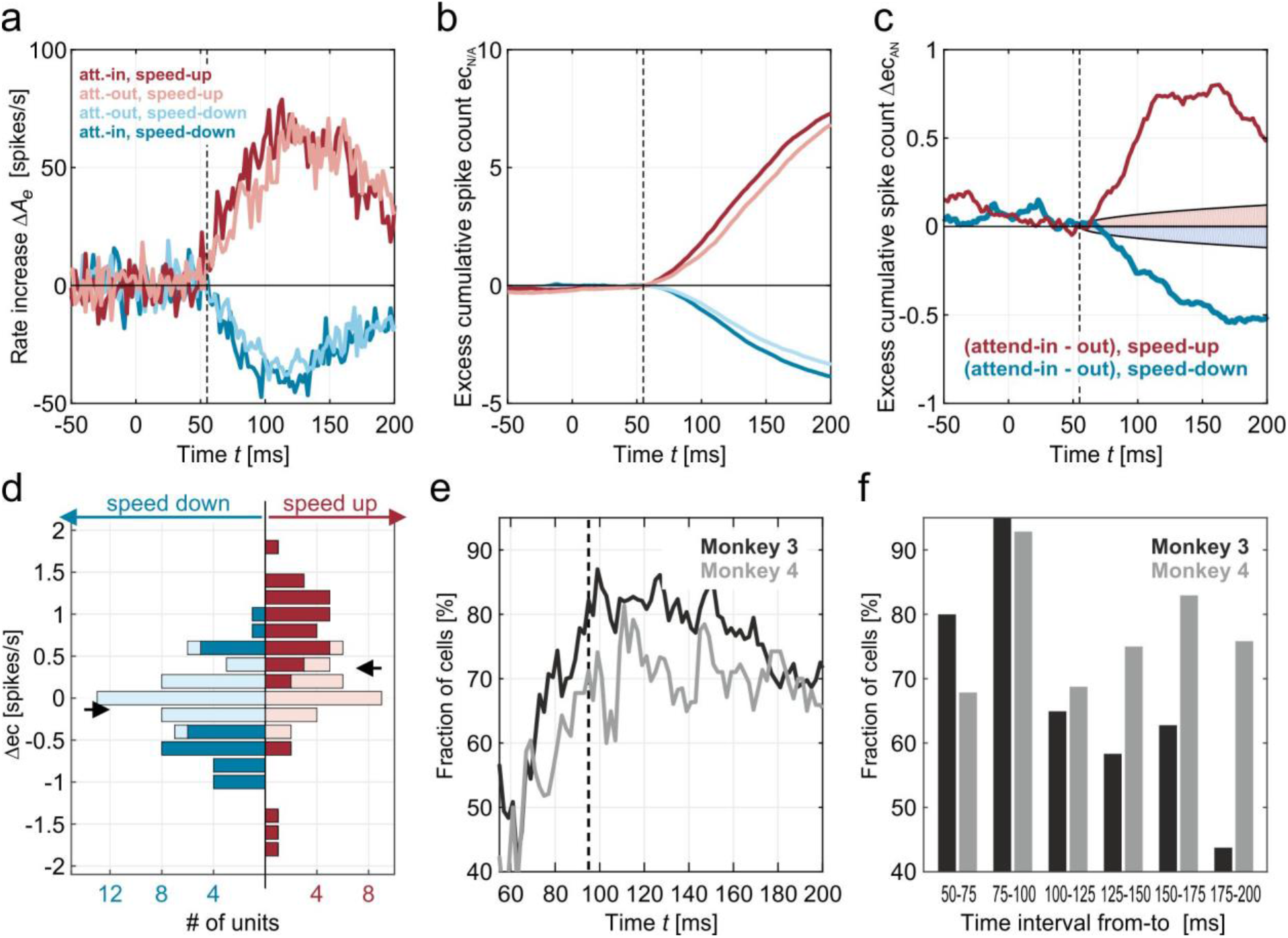
Quantification of attention-induced rate changes. **(a)** Population average of speed change-induced rate modulations for the four attentional conditions. Data taken from M3. **(b)** Corresponding excess cumulative spike counts relative to the pre-change firing rate. **(c)** Attention-induced differences in excess cumulative spike count, separately for accelerations (red) and decelerations (blue). Differences are significantly larger than 0 (*p* < 0.01) outside shaded regions. **(d)** Histogram of attention-induced differences in excess cumulative spike count over individual sites, separately for acceleration and deceleration conditions. Dark colors indicate samples with values being significantly different from zero. Samples were taken at 95 ms after stimulus change. **(e)** Percentage of significantly modulated individual units for which cumulative counts were significantly larger with attention in the speed-up condition or significantly smaller with attention in the speed-down condition. Black line, M3; gray line, M4. Dashed line indicates the time for which the samples for the histogram in panel (d) were taken. **(f)** Percentage of significantly modulated units for which spike count differences between activities before and after the stimulus change were significantly larger in the speed-up condition or significantly lower in the speed-down condition, separately for successive time intervals of 25 ms length. Black bars, M3; gray bars, M4.

Second, for analyzing the transient’s peak in response to the stimulus change, the firing rate was calculated in consecutive bins of 25 ms width. For both speed increments and decrements, up to more than 90 % of units were found to have significantly higher and lower rates, respectively, in the attend-in condition during the time intervals of 50 to 100 ms post-change (Fig. 6f). Yet, because this ratio decreased rapidly for later bins of the transient, consistent attentional modulation was limited to a brief period immediately following the stimulus change.

Taken together, the experimental data confirmed the model predictions for both the effect of attention onto the slope of positive and negative transients and, for the initial part of the transient, onto the modulation of relative peak responses for time constant ratios of *τ_i_/τ_e_* ≫ 1, as found in the response onset fits (Fig. 2e). As a novel physiological result, they provide evidence that attention modulates the same features of a negative transient than it does for positive transients, suggesting that processing of visual information, and its perception, can rely on information contained in reductions of firing rates.

## Discussion

The ability to detect rapid changes in complex, ever-changing environments is fundamental for both animal and human behavior. Neuronal responses to fast stimulus transitions usually come as brief episodes of increased or decreased neuronal activity, followed by a steady-state level of lower absolute amplitude. Pronounced transient changes in neuronal activation were observed in the brain of many different species, spanning the range from invertebrates to primates^23–25^, suggesting that they represent a basic principle in neuronal network dynamics. We here show that such a canonical computation can be realized by a very simple circuitry, essentially built of only one excitatory and one inhibitory unit, in which the excitatory unit’s output time course is normalized through divisive inhibition. The circuitry can be expressed by a set of equations with only three free parameters, obtained by fitting the model to each neuron’s onset response (to cover the individual unit’s kinetics). The simplicity of the model allowed for the comprehensive formal analysis of neuronal response dynamics and to reproduce and predict physiological transients to a significant range of stimulus transitions, including the interesting case of attentional modulation of rate-decreasing events.

### Non-stationary normalization by divisive inhibition

A key element of the model is normalization of the circuitry’s output by divisive inhibition. Normalization by divisive (shunting) inhibition was initially suggested as a means to explain nonlinearities in the response of neurons in visual cortex^17^. It consists of dividing the response of a given neuron, or group of neurons, by the average response of a pool of normalizing units, either within the same cortical area or between areas^26–30^. Since its introduction, the concept was successfully used to explain neural response characteristics in a range of different contexts and neural systems, both with and without attention (cf. Refs. [18],[31] for overview). Most work, however, has focused on static divisive normalization to describe modulations of sustained neuronal responses, while the circuitry we here introduce explicitly addresses the temporal response dynamics. Static normalization in our model emerges as the fixed point of the activation dynamics for a constant input. Static models may be capable to reproduce MT responses to dynamic stimuli to some extent^32^, but they have limits to capture fast neural responses and they are inappropriate to explain nonmonotonic transient responses to sudden input changes. Temporal low-pass filtering was previously suggested to circumvent these limitations and allowed modelling the time course of MT responses during pursuit eye movements^33^. Because low-pass filtering converts an instant input change to an exponentially saturating input current, divisive inhibition is delayed with respect to excitation and thus enabled generation of transient responses that successfully fitted the experimental data. The relaxation differential equations in our model have exactly the same effect – low-pass filtering the input drive, and delaying inhibition by assuming a larger time constant for the rise of the divisive term. However, the earlier model has a large number of free parameters and multiple functional modules for allowing detailed fits of neural responses to pursuit eye movements of different velocity^33^, while the model we here introduce has its focus on structural simplicity. This property enables thorough formal analysis for a large variety of stimulus conditions, with and without the effects induced by attention, and allowed to predict neuronal response dynamics under so far untested experimental conditions.

### Relation between transients and information processing, perception, and behavior

Biologically relevant signals usually occur on very short time scales, and the brains of both vertebrates and invertebrates generate rapid visual percepts, decisions, and motor behaviors. Flies, for instance, track and chase other flies with response times as short as 30 ms^34^, carnivorous vertebrates possess extremely fast sensorimotor programs for visually guided pursuit predation^35^, and primates, both human and nonhuman, categorize objects and perform appropriate motor responses within tens of milliseconds^36–38^. These findings imply that, while different behaviors in different species may involve different neuronal substrates and mechanisms, the brain strongly relies on fast neuronal codes to account for the strong temporal variability of sensory input during eye movements, self- and object-motion. Transient firing rate changes of small groups of neurons in response to sudden changes in sensory input are likely part of this code. Accordingly, such transients not only participate in detection of objects and events but carry detailed information about stimulus properties. In monkey temporal cortex, transients were shown to exhibit specificity for different head views within 25 ms following onset of the population response, and to contain more information than later epochs of the response^39,40^. A corresponding pattern was found in primary visual cortex V1, reaching a peak for detectability and discriminability of oriented gratings within 150 ms of the onset response, and in area MT, where most information on motion direction is available within the first 100 to 200 ms after stimulus onset^3,4,41^. This higher information content of onset transients is likely due to a larger gain and smaller variance as compared to steady-state activation levels during continuous stimulation^41,42^. Accordingly, because brief episodes of coherent motion or rapid speed changes were found to induce firing rate changes significantly correlating with behavioral choices, transients were linked to perceptual judgments^9,43–45^. The results of the current study show that the simple circuitry we used to implement rapid firing changes in response to input changes is fully reproducing the experimentally observed MT responses, including the over- and undershooting during change transients in comparison to firing rate changes expected from steady-state tuning properites^7^. Because thresholding of transients allowed for read-out of information in full accordance with human behavioral performance^6^, both for rate increments and decrements^7^, they provide a mechanistic explanation for computations within sensory cortex underlying the perceptual process of change detection and discrimination, basically realizable by non-stationary divisive inhibition within a simple cortical circuitry.

### Modulation of change transients by selective attention

Due to the close relation between transient firing rate patterns and perceptual judgments on the one hand, and the influence of selective attention on neuronal processing and behavioral performance on the other, transients are likely targets for attentional modulation. Recent monkey neurophysiological studies reported attention-dependent modulation of amplitudes, latencies, and gamma coherence during stimulus onset or stimulus change responses in various visual areas^9,19,45–49^, and a close relation between reaction times to attended speed increments and the latency of the change transient^9^. All of these results, however, were obtained by investigating stimulus events eliciting an increase in firing rate. If specific parameters of transient firing rate changes indeed underlie perceptual performance, the question arises how attention influences a population of neurons for stimuli inducing a decrease in mean activation. The formal analysis of our model dynamics predicted – not without surprise – that negative transients would basically be modulated by attention in the same way as positive ones, with the slope and the relative peak height being the response features to allow for consistent attentional modulation independent of the pre-change activation level of the neuron, and independent of the sign of the transient. The experimental confirmation of this prediction provides new physiological insights for understanding change detection and its modulation by attention. As an important result, they indicate a relevant constrain of task-dependent modulation. Because attention was implemented as a positive gain to the input of the circuitry in the model, firing rates during the pre-change epoch were always larger with attention than without, regardless of the type of change occurring later. The new physiological results reported here show that, albeit speed increments and decrements were presented block-wise and allowed the animals to make correct predictions on the sign of the upcoming change, attention consistently increased neuronal responses during the pre-change epoch by about the same factor. These results strongly suggest that attention-dependent modulation in early visual cortex is generally associated with a positive gain of neuronal responses, as opposed to a mechanism modulating responses in the same direction as the sensory event. Moreover, regardless of an always positive gain modulation during the pre-change period, the model predicted steeper response slopes with attention for both stimulus changes inducing an increase and a decrease in neuronal firing. This prediction was a direct consequence of the model’s inherent dynamics, since apart from input gain no other parameter of the model was changed to implement attention. In line with this prediction, the physiological experiments show a significant influence of spatial attention on both the rise and the decay time of the change transient, being steeper with attention than without, i.e. attentional modulation was independent of whether the stimuli induced an increase or a decrease in the firing of neurons. Based on the model’s temporal dynamics, a mechanistic explanation of this effect is that the stronger drive of both the excitatory and inhibitory unit allows a faster effect of divisive normalization with attention. Because normalization is acting in the direction of the input change, it is affecting the slope of both positive and negative transients likewise. Both the computational and the physiological data suggests that not only rapid positive changes in firing rate, but also rapid negative changes provide important information to downstream areas that are used for subsequent visual processing.

## Methods

### Electrophysiological data

MT data used to develop the model were recorded in the context of previously published studies^7,10^. Additional data to test model predictions were recorded using non-human primate standard behavioral and neurophysiological procedures. Housing of animals, experimental and surgical procedures were all in accordance with the *Directive 2010/63* issued by the European Commission and the *Regulation for the Welfare of Experimental Animals* issued by the Federal Government of Germany, and were approved by the local authorities. Data were acquired from two male macaque monkeys, six and eight years old. Recordings were performed using tungsten microelectrodes (2–5 MOhm, 125 mm shank diameter; Frederic Haer, Bowdoin, ME) and standard electrophysiological equipment. The pre-amplified signal was filtered between 0.7 and 5 KHz and sampled with a frequency of 25 KHz. Spikes were detected online by thresholding the signal. All spike data were then subjected to offline semiautomatic spike sorting using Klustakwik^50^, followed by manual adjustment of spike clusters using a custom-made algorithm for spike form and spike parameter illustration^51^. Visual stimulation, control, and documentation of behavioral data was performed using custom-made Matlab scripts and in-house software. Eye movements were controlled by a custom-made video-oculography system with 0.2 degree resolution.

### Visual stimulation and behavioral paradigm

Monkeys were tested in a behavioral task requiring detection of a rapid change in the speed of a moving stimulus, either an acceleration (speed change by a factor of ~2), or a deceleration (speed change by a factor of ~0.5), presented block-wise. The basic task design was the same as in previous studies^9,10^. Each trial started with appearance of a small red fixation spot (0.14 degree side length) at the center of the screen (22 inch cathode ray tube monitor, resolution 1,280 x 1,024 pixel, 100 Hz refresh rate). Monkeys initiated the trial by gazing at the fixation point, pressing a lever, and keeping it hold. Following a delay of 250 ms, a spatial cue appeared for 700 ms to indicate the location of the upcoming target stimulus, followed by another delay of 200 ms and subsequent appearance of two static Gabor stimuli (sine wave spatial frequency: 2 cycles/degree, Gaussian envelope: σ = 0.75 degree at half height), one centered above the RF of the recorded neuron and the other one mirrored across the fixation spot. Mean Gabor luminance was identical to background luminance (10 cd/m^2^). Gabors started to intrinsically move (speed: 2.17 degree/sec) 200 ms after onset, with motion direction adjusted to the preferred direction of the recoded neuron, as described elsewhere^9^. In about 40% – 50% of the trials, the uncued stimulus changed speed before the target stimulus, which had to be ignored by the monkeys. Following the speed change of the target, monkeys had to keep fixation for another 300 ms (to avoid contamination of the neuronal post-change response by eye movements) and to release the lever within a response window of 150-750 ms. Speed changes occurred within 0.66 and 5.5 sec after motion onset. Subsequent trials were separated by an intertrial interval of 3 – 4 sec. Releasing the lever outside the response window and eye movements of more than one degree from the fixation point caused immediate termination of a trial. Monkeys were rewarded with a few drops of water or diluted fruit juice for each correctly performed trial.

### Model steady-state activation and response to step functions

The model consists of two units driven by external input *I*, with one unit providing divisive inhibition on the other unit (Fig. 1b). Their dynamics are described by the differential equations (1)(2) and threshold-linear gain functions (3)-(4), with the steady-state activation resulting from a constant, suprathreshold input *I*_0_ given by

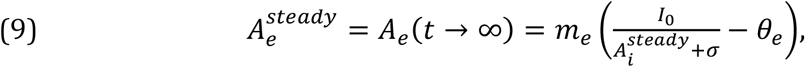

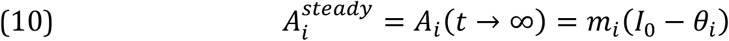

Inserting (10) into (9) provides equation (5). Assuming zero thresholds *θ_e_* = *θ_i_*, = 0 and a suprathreshold input *I*(*t*) > 0 allows to replace (3)-(4) by linear gain functions, simplifying further analysis. Instantaneous stimulus changes are considered as step functions providing a change in constant external input from *I^pre^* to *I^post^* at *t = t_change_*. Assuming that the model is in its steady state for *t < t_change_*, equation (2) can be explicitly solved and equation (1) for post-change activation (*t* ≥ *t_change_*) can be written as:

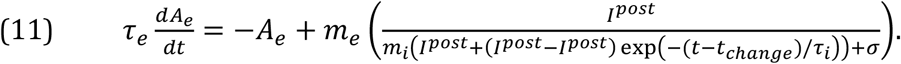

Using equation (5), *I^pre^* and *I^post^*, which are unknown in an experiment, can be expressed as functions of their corresponding steady-state (sustained) activations 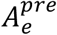 and 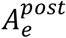 via equation (6).

### Model fit to physiological data

88 recordings from M1 and 54 recordings from M2 (single- and multi-units) were used to fit the model to stimulus onset transients. Transients were caused by the appearance of a grating inside a neuron’s RF, moving into its preferred direction while monkeys performed a simple fixation paradigm (Ref. [7] for experimental details). While the model responds immediately to a change in its input at time *t_change_*, the physiological change response is delayed by the processing time between retina and area MT. To account for this, response onset delay *Δτ* relative to stimulus onset was estimated by visual inspection of the PSTHs binned at different temporal resolutions, individually for each unit (averages: M1: 31ms +/− 12ms SD, M2: 32 +/− 10ms). Sustained responses 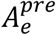 and 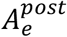 before and after stimulus change were determined by computing the average spike rate over all trials in the intervals [-100ms, 0ms] and [200ms, 500ms], respectively (time denoted relative to stimulus onset). 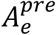 and 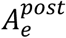 allow to numerically solve equation (6) and compare it directly to the delay-compensated, trial-averaged physiological response by computing the quadratic error 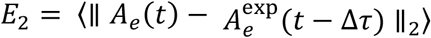, averaged over a 200ms interval after response onset. Experimental 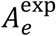 was sampled within 5 ms time bins and the model’s response was downsampled to the same temporal resolution for comparison. Parameters *τ_e_, τ_i_*, and 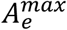 for explaining physiological dynamics were determined by an iterative grid search for minimizing *E*_2_. Because 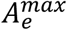 must be at least the value of 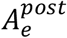, and values higher than 3 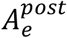 were never reached by the optimization procedure, initial grid search ranges for 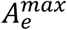 were set to [1.03 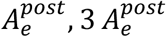]. Ranges for *τ_e_* and *τ_i_* were set to [1 ms, 100 ms] and [1 ms, 500 ms], respectively, with the upper limits being well above the observed rise/decay times of the PSTHs. The resolution of the search grid was 40 bins for 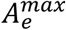, and 15 bins for the two time constants. Four iterations with subsequently refined grids resulted in a precision of 0.01 % for the parameter estimates within the initially chosen range. Goodness-of-fit was determined as the ratio 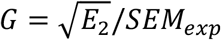 of the error 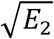 between model and trialaveraged physiological responses used in the fit (see above) to the standard error of the mean *SEM_exp_* (averaged over the time interval used for fitting).

### Calculation of transient and sustained activation changes

Under the condition *τ_e_* ≪ *τ_i_* (separation of time scales), the peak response during the transient can be approximated by:

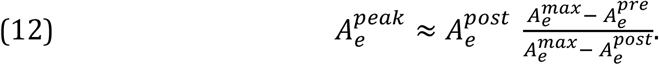

Under assumption of a log-Gaussian velocity tuning^14,15^, sustained activation can be expressed in terms of the stimulus velocity *v*:

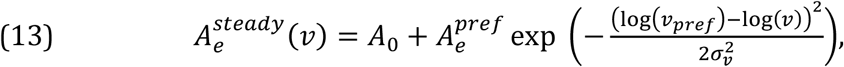

with spontaneous activity *A*_0_, tuning width *σ_v_* and maximum response amplitude 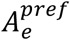 for preferred speed *v_pref_*. Computing 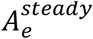 for stimulus velocities *v^pre^* and *v^post^* before and after the speed change, respectively, provides 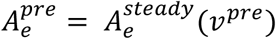 and 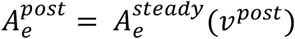 for equation (14), to obtain analytical expressions for 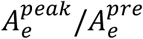 and 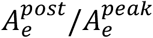 for arbitrary acceleration/deceleration ratios *v^pre^/v^post^* (Fig. 3). For evaluating 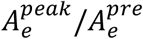 and 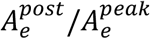 in comparison to experimental data (Fig. 3), we chose *A*_0_ = 0, a tuning half-width of 2.5 octaves (*σ_v_* = log(2^2.5^) ≈ 1.73) and 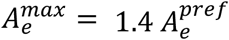.

### Modulation by attention

Attention is modeled by multiplicative modulation of input *I* by a factor *α* > 1, *I* → *αI* (Fig. 4a). Expressing absolute activation *A_e_* relative to 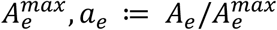 provides scaled pre- and post-change sustained activities 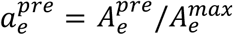 and 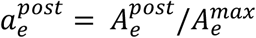 ranging in the interval [0, 1]. The initial slope 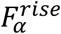 of the transient response as a function of 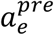 and 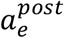 in dependence on *α* yields:

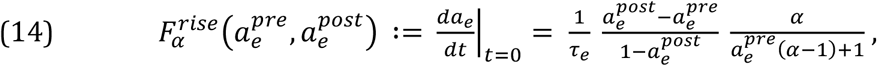

and the change in sustained activation level 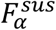 relative to the pre-change level becomes:

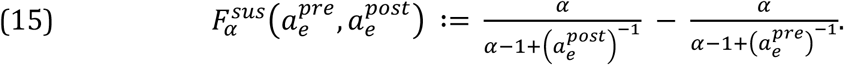

For assessing the attention-induced modulation in these quantities (Fig. 4b), we computed 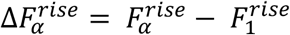 and 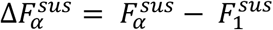. Peak activation 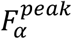 during transient responses must be computed numerically, thus 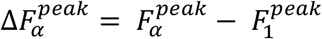 was obtained by solving equation (6) explicitly in dependence on the chosen time constants *τ_e_* and *τ_i_*.

### Calculation and comparison of physiological response parameters

In total, *N* = 45 sites were recorded in M3 and *N* = 25 sites in M4. For inclusion to data analysis, we required the speed-up condition to be associated with a significant firing rate increase in the attend-out condition (assessed in the time interval 140 to 160 ms after stimulus change, one-tailed test on Poissonian distribution around mean firing rate before stimulus change, p<0.05), which was given for *N_up_* = 36 sites in M3 and *N_up_* = 19 sites in M4. Similarly, the speed-down condition was required to be associated with a significant firing rate decrease in the attend-out condition, which was fulfilled for *N_down_* = 42 sites in M3 and *N_down_* = 21 sites in M4. Rise times and relative spike counts of the experimentally observed transients (Fig. 5) were calculated to compare physiological data against model predictions for Δ*F^rise^* and Δ*F^peak^*. First, rise times of physiological transients were assessed by determining excess cumulative spike counts following the stimulus change, defined as the number of spikes exceeding the spike count of a neuron continuing to fire with its observed pre-change rate. By visual inspection, we estimated an average response delay of *t_change_* = 55 ms for both monkeys. For estimating pre-change activity, we computed the summed firing rate *F^pre^* over all trials of a given attention condition in the time window [-400ms, *t_change_*]. Excess cumulative spike count *ec*(*t*) was then defined for *t* > *t_change_* by:

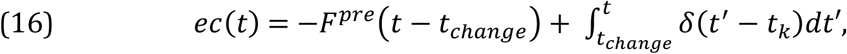

where *t_k_* denote the times of *K* spikes *k = 1, …, K*, considering all trials of the respective condition. For comparing the initial slopes of two attention conditions *N* and *A*, we first computed their cumulative excess responses *ec_N_*(*t*) and *ec_A_*(*t*). We then evaluated the difference Δ*ec_AN_*(*t*) = *ec_A_*(*t*) – *ec_N_*(*t*) and tested statistically whether Δ*ec_AN_*(*t*) was significantly deviating from zero. The transient of condition *A* was considered to rise significantly faster (or slower) than the transient of condition *N*, if Δ*ec_AN_*(*t*) was above (or below) a time-dependent threshold. Thresholds were set to 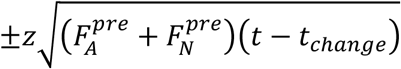, where *z* = 2.32 was chosen to yield a significance level of *p* < 0.01.

Second, spike counts for individual attention conditions and during different periods of the transient were calculated as the difference Δ*ac* between the absolute spike count before and after the speed change, Δ*ac*(*itv*) = *ac^post^*(*itv*) — *ac^pre^*, where *itv* denotes intervals of 25 ms length, taken between 50 ms and 200 ms following the speed change for the transient. The steady-state response before the change was estimated during the period from −400 ms to *t_onset_*. The difference between two relative spike counts was considered significantly different from zero when it exceeded 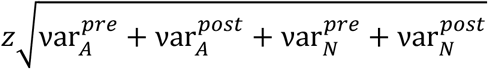, where var designates the spike count variance in the corresponding attentional condition pre- or post-stimulus change.

## Acknowledgments

The authors thank Franziska Müller, Peter Bujotzek, Katrin Thoss and Ramazani Hakizimani for technical and training support, Bonne Habekost for helping to re-organize physiological data, and Andreas Traschütz for his insightful work on visual transients. The work was supported by BMBF grant 01GQ1106 and DFG grant WE 5469/2-1 and 3-1.

## Author Contributions

U.E. and D.W. conceived and designed research, U.E., X.C., and L.B. performed computational work and experiments, F.O.G. performed electrophysiological experiments, U.E. and D.W. interpreted results of experiments, U.E. and D.W. prepared figures, U.E. and D.W. wrote manuscript, U.E., X.C., L.B., F.O.G., and D.W. approved final version of manuscript.

